# Time affects a time-consuming, flexible component and a fast, inflexible component of sensorimotor adaptation

**DOI:** 10.1101/2024.11.17.624025

**Authors:** Li-Ann Leow, Welber Marinovic, Scott Albert, Timothy J Carroll

**Affiliations:** School of Arts and Humanities, Edith Cowan University, Joondalup, Western Australia, Australia; School of Psychology, The University of Queensland, St Lucia, Queensland, Australia; Centre for Sensorimotor Performance, School of Human Movement & Nutrition Sciences, St Lucia, Queensland, Australia; Curtin School of Population Health, Bentley, Western Australia, Australia; University of North Carolina Chapel Hill, United States of America

## Abstract

Prior learning can impair future learning when the requirements of the two memories conflict, a phenomenon termed anterograde interference. In sensorimotor adaptation, the passage of time between initial and future learning can reduce such interference effects. However, we still do not fully understand how time affects learning, as some studies found no effects of time on interference. One possible explanation for such inconclusive findings is that time affects distinct processes underpinning sensorimotor adaptation differently, and these processes may compensate for each other’s effects on behaviour. Here, we used task manipulations that (1) dissociate adaptation processes driven by task errors from adaptation processes driven by sensory prediction errors, and (2) separate the task-error driven adaptation processes into a flexible component that could not be acquired under time-pressure, or a less-flexible component that could not be acquired under time-pressure. The time between initial and subsequent learning seemed to alter both the flexible and inflexible components of adaptation driven by task errors. Time also led to a small reduction of interference arising from sensory prediction errors. Thus, we provide evidence that multiple components of sensorimotor adaptation are sensitive to the passage of time.

---

An Australian tourist driving a left-hand-drive car in North America is prone to errors, such as activating the windscreen wipers instead of indicating, and driving onto the wrong side of the road after making a turn. Such impairment of new learning as a result of interference from prior learning is termed proactive or anterograde interference (Underwood, 1948). Anterograde interference is common to multiple forms of learning across cognitive, motor, and perceptual learning domains. For example, interference is evident when successively adapting to opposing perturbations of sensory feedback (e.g., Flook & McGonigle, 1977; Brashers-Krug *et al*., 1996; Wigmore *et al*., 2002; Sing & Smith, 2010; Kumar *et al*., 2018; Lerner *et al*., 2020), and when successively learning two different perceptual learning tasks (Seitz *et al*., 2005). Interference can even occur between different forms of motor learning that require different movement kinematics (Krakauer *et al*., 1999; Tong *et al*., 2002). Effects can be long lasting, persisting up to 5 months (Shadmehr & Brashers-Krug, 1997) or a year (Yamamoto *et al*., 2009). Yet, despite the ubiquity and persistence of anterograde interference, in the domain of motor learning, the mechanisms that underlie the effect remains incompletely understood.

An important determinant of anterograde interference effects is the passage of time after initial learning. A large body of work showed that time systematically reduces the size of anterograde interference effects, such that a period of ≈6 hours reduced anterograde interference (Brashers-Krug *et al*., 1996; Shadmehr & Brashers-Krug, 1997; Krakauer *et al*., 2005; Overduin *et al*., 2006; Lerner *et al*., 2020; Albert *et al*., 2022b; Solano *et al*., 2024). However, the effect of time on interference is somewhat controversial, as some studies prominently found no evidence that the passage of time reduces anterograde interference (e.g., Bock *et al*., 2001; Goedert & Willingham, 2002; Caithness *et al*., 2004; Miall *et al*., 2004). One idea posited to explain this discrepancy in findings was that that non-specific meta-learning effects (i.e., “learning-to-learn” effects that cause general improvements in learning) masked interference effects, which rendered interference effects less sensitive to the effects of time (Miall *et al*., 2004). Another possible explanation is that the effects of time were masked by retrieval inhibition effects, where previously consolidated memories are not readily retrieved because there is no cue to retrieve the appropriate motor output, such that a recency effect drives recall of the most recent learning (Krakauer & Shadmehr, 2006; Krakauer, 2009). This explanation is supported by findings that returning behaviour to the un-adapted state via no-perturbation “washout” trials reinstates consolidating effects of time on motor memories (Krakauer *et al*., 2005). However, the explanation does not readily explain the absence of the effects of time in conditions with washout (Caithness *et al*., 2004), and the finding that washout itself causes interference effects (Hinder *et al*., 2007). Thus, we still have an incomplete understanding of how time affects motor memories in sensorimotor adaptation.

Another possible explanation is that time has distinct effect on the multiple components of adaptation, which can have mutually compensatory effects (Hwang *et al*., 2006; Mazzoni & Krakauer, 2006; Taylor *et al*., 2014; Miyamoto *et al*., 2020; Albert *et al*., 2022b), and so could mask the effects of time on net behaviour. These components of learning can be experimentally dissociated by considering the distinct processes that drive them (Izawa & Shadmehr, 2011; Leow *et al*., 2018; Tsay *et al*., 2022). For example, sensory prediction errors, or discrepancies between predicted and actual sensory outcomes of movement, appear to drive an automatic, obligatory updating of the mapping between motor commands and predicted sensory outcomes (Mazzoni & Krakauer, 2006). Task errors, or discrepancies between predicted and actual task outcomes, might not only drive a person to devise a deliberate strategy to compensate for the perturbation (Hwang *et al*., 2006), but also trigger the formation of automatic, inflexible associations between aspects of the movement and the task context (McDougle & Taylor, 2019; Leow *et al*., 2020). Here, we asked if the memories that underlie the different components of adaptation driven by task errors and sensory prediction errors might respond differently to the passage of time, and thus have different effects on adaptation to a subsequent task with conflicting requirements. We experimentally teased apart the multiple components of adaptation and found that the effect of learning driven by task errors on subsequent adaptation to visuomotor rotation was sensitive to the passage of time. Time enhanced interference from the fast and inflexible components of adaptation to task errors on subsequent visuomotor adaptation but caused a paradoxical enhancement in visuomotor adaptation for the time-consuming and flexible component of learning. By contrast, there was a small decay in the interference from learning driven by sensory prediction errors on subsequent adaptation to opposing visuomotor rotation.

## Materials & Methods

### Participants

Eighty-eight participants (25 males, age range 17-34 years, mean age 20.5) completed Experiment 1. Forty-three participants completed Experiment 2. All participants were naïve to visuomotor rotation and force-field adaptation tasks, and were naïve to the aims of the study. Participants received course credit or monetary reimbursement upon study completion ($20 per hour). The study was approved by the Human Research Ethics Committee at The University of Queensland. All participants provided written informed consent. This study conforms to the Declaration of Helsinki.

### Apparatus

Participants completed the task using a vBOT planar robotic manipulandum, which has a low-mass, two-link carbon fibre arm and measures position with optical encoders sampled at 1000 Hz (Howard et al., 2009). Participants were seated on a height-adjustable chair at their ideal height for viewing the screen for the duration of the experiment. Visual feedback was presented on a horizontal plane on a 27” LCD computer monitor (ASUS, VG278H, set at 60Hz refresh rate) mounted above the vBOT and projected to the participant via a mirror in a darkened room, preventing direct vision of their hand. The mirror allowed the visual feedback of the targets, the start circle, and hand cursor to be presented in the plane of movement, with a black background. The start was aligned approximately 10cm to the right of the participant’s mid-sagittal plane at approximately mid-sternum level. An air-sled was used to support the weight of participants’ right forearms, to reduce possible effects of fatigue.

### General Trial Structure

Using their right hand, participants held the robotic arm, whose endpoint position was represented on-screen as a 0.5cm radius red circle, which appeared when the hand came within 1 cm of the central start position (a 0.5cm white circle). In each trial, the central start circle first appeared, and participants were instructed to move the cursor to the start circle. If participants failed to move their hand to within 1cm of the start circle after 1 second, the robotic manipulandum passively moved the participant’s hand to the start circle via a simulated 2-dimensional spring with the spring constant magnitude increasing linearly over time. A trial was initiated when the cursor remained within the start circle at a speed below 0.1 cm/s for 200ms. Then, targets (0.5cm radius yellow circles) appeared in random order at one of eight locations (0°, 45°…. 315°), 9 cm away from the start circle. Participants were instructed to move the cursor through the target in a slicing action. Across all trials (except the no-feedback trials, described later) cursor feedback was only provided after the cursor had travelled more than 4 cm of the 9 cm movement, and terminated immediately upon attaining 9 cm movement extent. After completing every trial, cursor feedback was provided again when the cursor came within 1 cm of the start.

### Time pressure manipulations

Similar to our previous work (Leow *et al*., 2017), we used a timed-response paradigm (e.g., Schouten & Bekker, 1967) to enforce long movement preparation times (1000ms) for all trials in Experiment 1 & 2, except for the training and washout block in Experiment 2, which had short movement preparation times of less than 200ms. Specifically, a sequence of three tones spaced 500ms apart was presented via external speakers, and participants were instructed to time the onset of the third tone. Targets appeared at 1000 ms (long preparation time trials) or 200 ms (short preparation time trials) minus a monitor display latency (27.6 ± 1.8 ms), before the third tone. Target direction information became available 972.4ms before the desired initiation time for long preparation time trials, and 172.4ms before the desired initiation time for short preparation time trials. Movement initiation was defined online as the time when hand speed exceeded 2cm/s. When movements were initiated more than 100 ms before the desired initiation time, the trial was aborted: the screen was blanked and a “Too Soon” on-screen error signal appeared, and no visual feedback about movements was available. When movements were initiated 50 ms later than the third tone, the trial was aborted: the screen was blanked and a “Too Late” on-screen error signal appeared. Thus, movements had to be initiated between 872 and 1022 ms of target presentation in Experiment 1 and between 122 and 272 ms of target presentation in Experiment 2.

### Experiment 1: Manipulating task errors at training and the passage of time after training

We used an A1-B design to quantify anterograde interference, where participants first adapt to one perturbation at initial adaptation (A1) and then adapt to a directionally opposite perturbations in block B. In addition, we used washout to return behaviour to the unadapted state after each perturbation block. This contrasts with standard anterograde interference study protocols which have participants adapt to successive opposing perturbations without washout, and provided a stronger test for interference effects that persist despite returning behaviour back to baseline.

In a 2 x 2 design, we manipulated task error history at initial learning and subsequent washout (no task error history denoted as **TE-**, task error history denoted as **TE+**), and the passage of time after initial learning and subsequent washout (no time delay denoted as **Time-**, time delay denoted as **Time+**). Participants were assigned to one of four conditions: **TE-Time-** (n=26), **TE-Time+**(n=20), **TE+Time-** (n=22), **TE+Time+** (n=20).

### Block structure

Participants first completed an unrecorded familiarization block (40 trials, or 6 bins) to familiarise themselves with the target reaching task, followed by a baseline block (40 trials, or 6 bins) with veridical cursor feedback. Next, in the A1 block (thereafter termed training) (30 bins, or 240 trials), a 30° cursor rotation was abruptly imposed (rotation direction was counterbalanced between participants), followed by a no-rotation washout block (A1 washout, 30 bins) to return behaviour to the unadapted state. Both blocks A1 and A1 washout were conducted according to the task error manipulation designated for each group.

### Time manipulations

Participants assigned to the time delay conditions then had an overnight delay, whilst their no-delay counterparts went straight to the next block. After the time delay manipulation, all participants encountered block B (30 bins, or 240 trials), with a 30° cursor rotation in the opposite direction to that at training. Block B contained the no-task-error manipulation, described below, for all groups.

Block B was followed by **B washout** with task errors (30 bins). For the TE-groups, as all preceding blocks were without target-errors, B washout was the first time they encountered task errors resulting from an abrupt change in cursor feedback. Finally, all participants completed a final A2 block with the same cursor rotation direction as A1, but with task errors for all groups. Results from this A2 block will be reported in a separate manuscript for parsimony.

After each adaptation block, participants were explicitly notified that the perturbation had been removed, and asked to move straight to the target in the subsequent bin of 8 **no-feedback** trials, where cursor feedback was hidden as soon as the cursor left the start. Persistently adapted behaviour despite explicit knowledge of perturbation removal (i.e., aftereffects) suggests an altered mapping between the outgoing motor command and its predicted sensory outcomes (Heuer & Hegele, 2008; Taylor *et al*., 2014) that cannot be volitionally disengaged, and thus appears implicit in nature.

### Task-error manipulations

At training (A1) and subsequent washout (A1 washout), participants in the TE+ conditions had each target remain stationary during each trial, and thus experienced typical task errors induced by the abrupt onset and removal of the cursor rotation. By contrast, participants in the TE-conditions were deprived of task errors in A1 and A1 washout. This was achieved by moving the target mid-movement to align with the cursor, analogous to moving a basketball hoop towards the ball mid-flight—the ball always goes through the hoop. Specifically, as soon as the cursor reached 4cm of the 9cm start-target distance, the target was extinguished and quickly re-appeared in the next frame in the same direction as the cursor. Note that no-cursor feedback was provided in the first 4cm of movement or beyond 9cm of the start-target distance to discourage online movement corrections. The first 4cm of cursor feedback was always unavailable across all trials in all conditions.

Between each block of trials, there was a small delay to allow loading of the different experimental blocks and/or experimental instructions. There were no structured rest breaks, but participants were allowed to take rest breaks between trials as needed. The cumulative test duration was approximately 2 hours.

### Experiment 2: Suppressing explicit strategies to reveal effects of an inflexible component

We previously showed that learning to counteract task errors improved subsequent adaptation to a novel cursor rotation that required a new movement solution, demonstrating flexibility of motor memories encoded by task errors (Leow *et al*., 2020). However, such flexibility disappeared under time-pressured conditions that suppressed the expression of explicit strategies (Leow *et al*., 2020). This suggests that task errors can induce multiple learning components with distinct features: a time-consuming but flexible component, and a time-efficient but inflexible component. In Experiment 2, we sought to understand whether time might also affect consolidation of the fast, inflexible learning component induced by task errors, by using time-pressure to suppress the expression of the time-consuming flexible learning component at initial adaptation. We used the same design as the task-error groups of experiment 1, except that a short-preparation-time manipulation was deployed to limit the time available to prepare movements (< 200ms) for the baseline, training (block A1), the no-feedback bin, and the first washout block (A1 washout). All other aspects of the task were the same as in Experiment 1. Participants were assigned to either experience an overnight time delay after A1 washout (**TimePressure+Time+ condition,** n=21), or to have no time delay after A1 washout (**TimePressure+Time-**, n=22). From block B onwards, all trials had the standard long-preparation time conditions. We compared these data to the task-error conditions in Experiment 1.

### Data analysis

Across all datasets, movement onset time was estimated as the time at which hand speed first exceeded 2 cm/s. Reach direction relative to the target direction was quantified at 20 percent of the movement distance. This procedure ensured that movement direction was quantified at less than 200 ms into the movement, at which time the size of online corrections in hand position is small (Elliott *et al*., 2001). Reach directions outside a +/- 90 degree range of the start-target direction were discarded from analyses (1.3% of all total trials). Similar to our previous work(Leow *et al*., 2020), the target-jump manipulations we employed to remove block B task errors resulted in more than 10% of block B reach directions outside a +/- 90 degree range in five out of the 129 participants (four of whom were in Experiment 2). Our outlier exclusion criteria excluded these reaches from the analyses. We also excluded one dataset from the TE+Time+ condition (clear circles in Figure D, red symbols), who showed aftereffects in the opposite direction.

For ease of interpretation, we calculated percent adaptation, by estimating adaptation as a percentage of reach directions relative to ideally adapted reach directions, similar to (Krakauer *et al*., 2005).

percent adaptation = 100% × (reach direction)/(ideal reach direction)

In Experiment 1, we aimed to quantify how the passage of time after initial adaptation and how a history of exposure to task errors during initial adaptation alters subsequent adaptation to a directionally opposite perturbation in B. In Experiment 2, we aimed to quantify how the passage of time after initial adaptation and how limiting preparation time during initial adaptation alters subsequent adaptation to a directionally opposite perturbation in B. To this end, we ran ANOVAs with between-subjects factors Task Error History (task error history, no task error history) and Time (time delay, no time-delay) for Experiment 1, and Preparation Time (short preparation time, long preparation time) and Time (time delay, no time-delay) for Experiment 2.

We were interested in the following dependent variables:

*Adaptation performance*, which was quantified by first binning trial-by-trial percent adaptation into bins of 8 trials, and then splitting the adaptation block into an early phase (first 10 bins), middle phase (second 10 bins) and late phase (final 10 bins) of each adaptation block. Then, repeated measures ANOVAs were run with the within-subjects factor block (A1, B), Phase (early, middle, late).

*Flexible learning*, which was quantified as the flexible change in reach behaviour upon verbal instruction that all perturbations have been removed and participants are now to move straight to the target. This change in reach behaviour upon instruction is typically interpreted as explicit strategy use (Hadjiosif & Krakauer, 2021; Maresch *et al*., 2021; Albert *et al*., 2022b), however, we and others have previously shown that such behaviour is not always strategic in nature, as time-pressure reveals a non-strategic facet to this learning, which can sometimes impair performance (McDougle & Taylor, 2019; Leow *et al*., 2020). As such flexible changes in behaviour upon instruction can occur rapidly across trials (Miyamoto *et al*., 2020), we used repeated-measures ANOVAs with within-subjects factors Instruction (Pre-Instruction, Post-Instruction) and Trial (1,…8) to quantify flexible changes in behaviour upon instruction. We chose to avoid estimating flexible learning as a change score from pre-to post-instruction, as change scores can be statistically inefficient (Vickers, 2001).

*Implicit aftereffects*, which were quantified as the magnitude of persistently adapted behaviour despite explicit knowledge of perturbation removal in the no-feedback trials. Aftereffects indicate an implicit remapping of the input-output relationship between the predicted and the actual sensory outcomes of movement that resists volitional disengagement, and are a measure of implicit learning. As aftereffects can decay across trials (Kitago *et al*., 2013), aftereffects were quantified via repeated-measures ANOVAs with within-subjects factors Trial (1…8).

### Statistical analyses

We report both Bayesian and frequentist statistics. Bayes factors do not have a tendency to over-estimate the evidence against the null hypothesis (Gelman & Tuerlinckx, 2000; Wetzels *et al*., 2011), and allows the evaluation of evidence for both the alternative hypothesis and for the null. Analyses were conducted in JASP (0.19.2). For frequentist statistics, an alpha-level of 0.05 was used, and cohen’s d values with 95% confidence intervals were used to quantify effect sizes. For Bayesian statistics, the default Cauchy prior widths (0.707) were used to quantify the relative evidence that the data came from the alternative versus a null model. For Bayesian ANOVAs, inclusion Bayes factors (*BF*inclusion, or BF_incl_) were determined to estimate the strength of evidence in favour of including a particular effect averaged across all candidate models that include the effect of interest (Wagenmakers *et al*., 2018), whereas exclusion Bayes factors (BFexclusion, BF_excl_) were used to estimate the strength of evidence against of including an effect averaged across all candidate models that exclude the effect of interest. For Bayesian t-tests, BF_10_ was taken as evidence supporting the test hypothesis, whereas BF_01_ was taken as evidence supporting the null hypothesis. For Bayesian statistics, Jeffreys’s evidence categories for interpretation (Wetzels *et al*., 2011) were used to evaluate reported Bayes Factors. Specifically, BF <1 were considered weak evidence for the alternative hypothesis, BF= 1-3 were considered as anecdotal to moderate evidence for the alternative hypothesis, BF of 3-10 were considered moderate to strong evidence for the alternative hypothesis, whilst BF of >10 was considered strong evidence for the alternative hypothesis.

## Results

### Experiment 1

#### Adaptation performance

Figure 2 top panels shows that for groups who initially trained with task errors, the passage of time after training improved subsequent adaptation performance when faced with a conflicting perturbation in block B (top), in contrast to the groups who initially trained without task errors, who appeared less sensitive to the effects of time (bottom), particularly in the late adaptation phase, as suggested by phase x task error history x time interaction, [BF_incl_ = 170.32, F(2,160) =2.17, p = 0.12, ω² = 0.002]. Simple effects analyses showed a beneficial effect of time during late phase adaptation performance for the task error groups (see Figure 2B) [moderate evidence for the effect of time, BF_incl_ = 6.45, F(1,80) = 8.154, p = 0.007], but not for the no-task-error groups [inequivocal evidence for the effect of time, BF_incl_ = 0.58, F(1,80) = 0.42, p = 0.52].

**Figure 1A:**
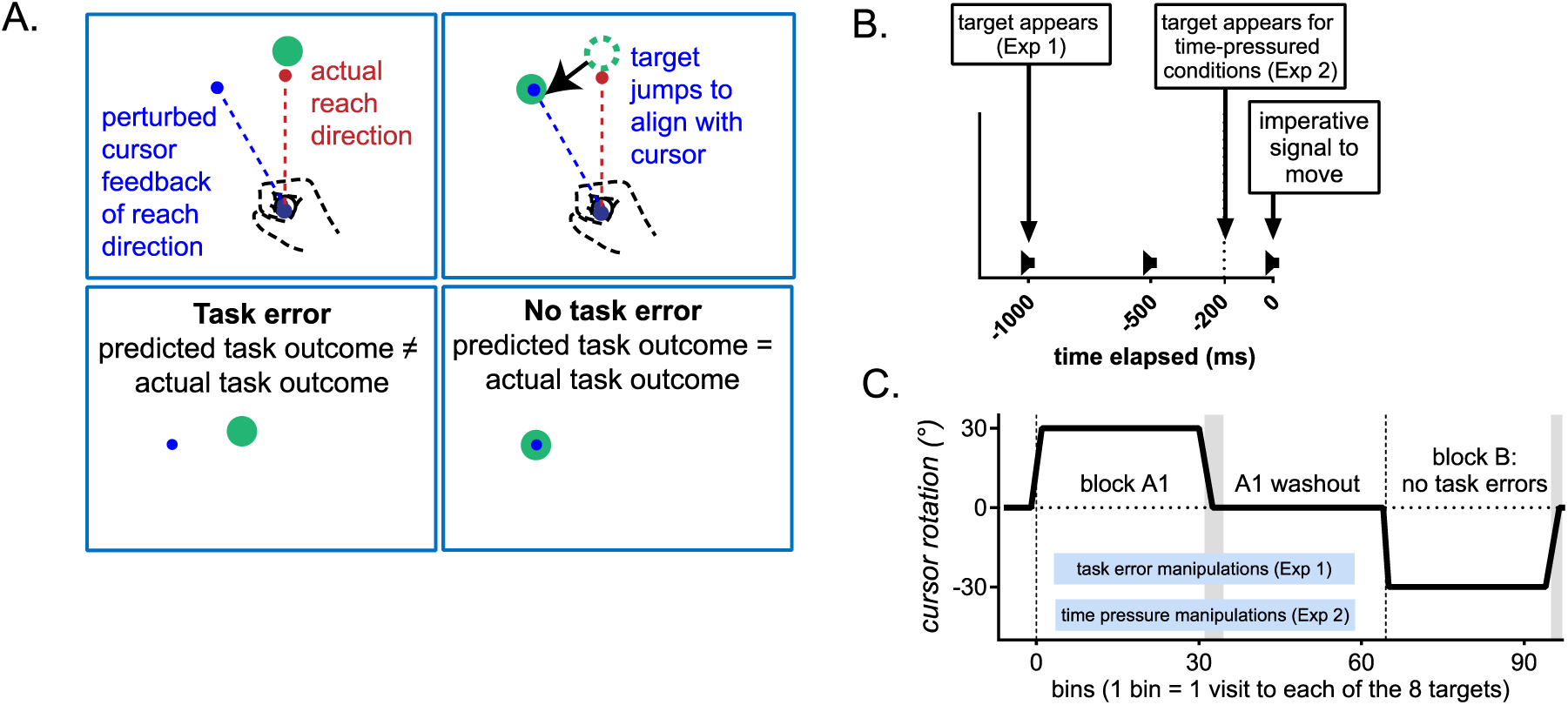
Task error manipulations. Under standard task error conditions, the perturbed cursor feedback caused the cursor to miss the target, eliciting a failure to achieve the instructed task goal of hitting the target (task error). Under no-task-error conditions, the target disappeared mid-movement (4cm into the 9cm start-target distance) and re-appeared within the next frame to align with on-screen cursor, such that the cursor perturbation did not elicit a failure to achieve the task goal of hitting the target. Figure 1B: The timed-response protocol which enforced long preparation times in Experiment 1, and time pressure (i.e., short-preparation times) in Experiment 2. Figure 1C. Task structure, where task error manipulations were applied at training and washout, whilst time delay manipulations were applied after washout. In Experiment 1, at training (A1) and subsequent washout, participants experienced task error manipulations (either no task error or standard task error). The Time+ groups were allowed an overnight delay, whilst the Time-groups proceeded directly to subsequent blocks. In Experiment 2, participants experienced time-pressure during training and washout. Rotation directions were counter-balanced in both experiments.

**Figure 2.**
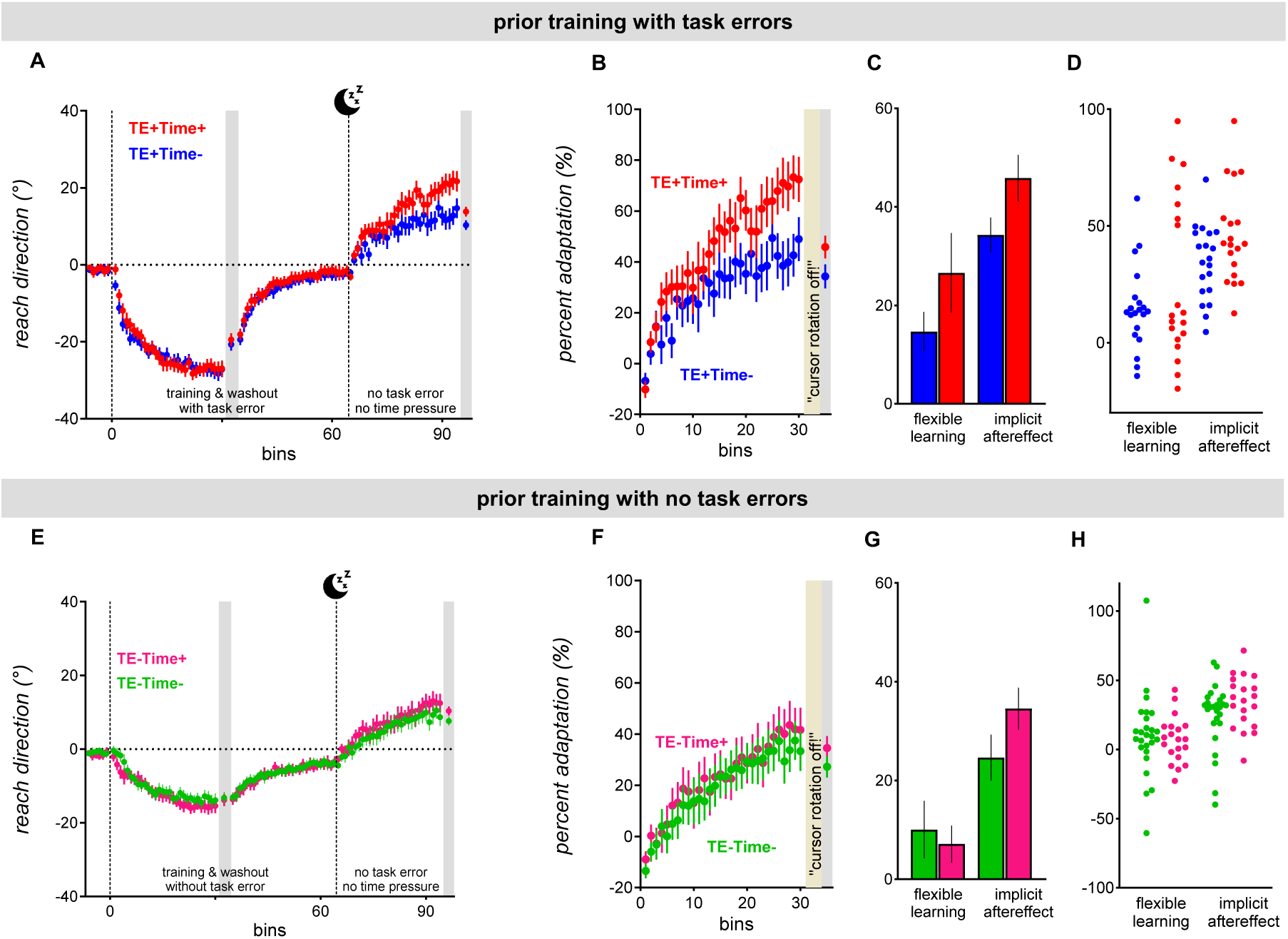
Adaptation in groups who completed prior training with task errors (top panel) or without task errors (bottom panel) in Experiment 1 (bottom panel), as evidenced in bin-averaged reach directions (A&E). Block B adaptation performance (B&F), and strategy use and aftereffects (C&D). Across all groups, the passage of time tended to increase implicit aftereffects (C&G). For those who trained with task errors, the passage of time after training improved subsequent block B adaptation performance (B) and flexible learning (C). Error bars are standard errors of the mean.

#### Flexible learning

Flexible learning, estimated via the volitional change in percent adaptation upon instruction, was more prominent in those with task error history, as shown by Instruction x Task Error History interaction, [BF_incl_ = 55.98, F(1,79) = 6.41, p = 0.01, ω² = 0.03]. Those who trained with task errors showed more flexible learning if they also experienced an overnight time delay after training [26.6%, 95%CI: [11.9%, 41.3%]), t(79) = 4.9, p_holm_ = 0.00003, Cohen’s d = 0.7, 95% CI [0.20, 1.19] than those experienced no overnight time delay [mean flexible learning: 14.7%, 95% CI: 0.35% 29.0%], t(79) = 2.77, p_holm_ = 0.03, Cohen’s d = 0.38, 95% CI [−0.07 0.84]. Those who trained without task errors showed relatively little flexible learning (average approximately 7%, both with the overnight delay (95% CI [−7.2%, 21.4%], cohen’s d = 0.19) and without the overnight delay (95%CI [−5.3%, 20.8%], (cohen’s d = 0.2, 95% CI [−0.21 0.61]). The passage of time did not alter flexible learning in those who trained without task errors, as shown in strong evidence for excluding the instruction x time interaction (BF_excl_ = 6.07) and for excluding the instruction x trial x time interaction (BFexcl = 29.83).

#### Implicit aftereffects

The passage of time after initial training reduced modestly reduced anterograde interference in implicit aftereffects resulting from adaptation to an opposing perturbation in block B, increasing the size of block B aftereffects, as shown by main effects of time [BF_incl_ = 2.69, F(1, 81) = 5.85, p = .02, ω² = 0.03], and post-hoc t-tests [BF_10, U_ = 1080.45, t(81) = 2.42, Cohen’s d= 0.33 [0.05 0.60], p_holm=_ 0.02]. Task error at training also modestly increased the size of block B implicit aftereffects, as shown by main effect of task error history [BF_incl_ = 2.50, F(1, 81) = 5.59, p = .02, ω² = 0.03], and post-hoc t-tests [BF_10, U_ = 725.81, t(81) = 2.36, Cohen’s d = 0.32 [0.05 0.59], p_holm=_ 0.02].

Thus, Experiment 1 showed that the passage of time after initial learning enhanced adaptation performance when subsequently faced with a conflicting perturbation, possibly by enhancing a flexible component of learning driven by task errors.

### Experiment 2

Previously, using time-pressure manipulations, we found evidence that task errors trigger a time-consuming component that improves subsequent adaptation to a conflicting perturbation, and a component expressible under time-pressure that can impair subsequent adaptation to a conflicting perturbation (Leow et al., 2020). In a similar vein, others have also demonstrated an adaptation component that is evident under time pressure (McDougle & Taylor, 2019). We wanted to test whether this component is sensitive to passage of time. Thus, in Experiment 2, we replicated the design of the task error groups, but additionally applied time-pressure in the initial adaptation block, as well as in the no-feedback block and the washout block that followed the initial adaptation block. We then tested how the passage of time after initial adaptation altered subsequent adaptation to an opposing perturbation in block B, where the expression of explicit strategies were disincentivised by removing task errors.

#### Adaptation performance

Prior training without time pressure improved adaptation performance when encountering an opposite perturbation in block B, despite the absence of task errors and time pressure in block B (see Figure 3B & F), as shown by strong evidence for a main effect of time-pressure, BF_incl_ = 22.29, F(1,76) = 9.98, p = 0.002, ω² = 0.06. The effect of training with time pressure depended on the passage of time after training [Time x Time-Pressure interaction, BF_incl_ = 2.02, F(1,76) = 4.49, p = 0.04, ω² = 0.02], particularly in the late adaptation phase (very strong evidence for the Time x Time-Pressure x Phase interaction, BF_incl_ = 4.12E+7, F(2,152) = 4.69, p = 0.01, ω² = 0.009]). In contrast to their counterparts who trained without time-pressure, whose late phase adaptation performance benefited from the passage of time after training (see Figure 3 red symbols), t(76) = 1.82, p_holm_ = 0.07, cohen’s d = 0.51 [−0.26 1.29]), the passage of time tended to *impair* performance in those who previously trained with time pressure (mean difference with time −26.9%, 95% CI [−51.6% −2.3%], t(76) = −2.17, p_holm_ = 0.03, cohen’s d = −0.6, 95% CI [−1.36 0.16]. Thus, the passage of time after training seemed to augment a component of adaptation acquired under time pressure, which impaired late phase adaptation performance during block B.

**Figure 3.**
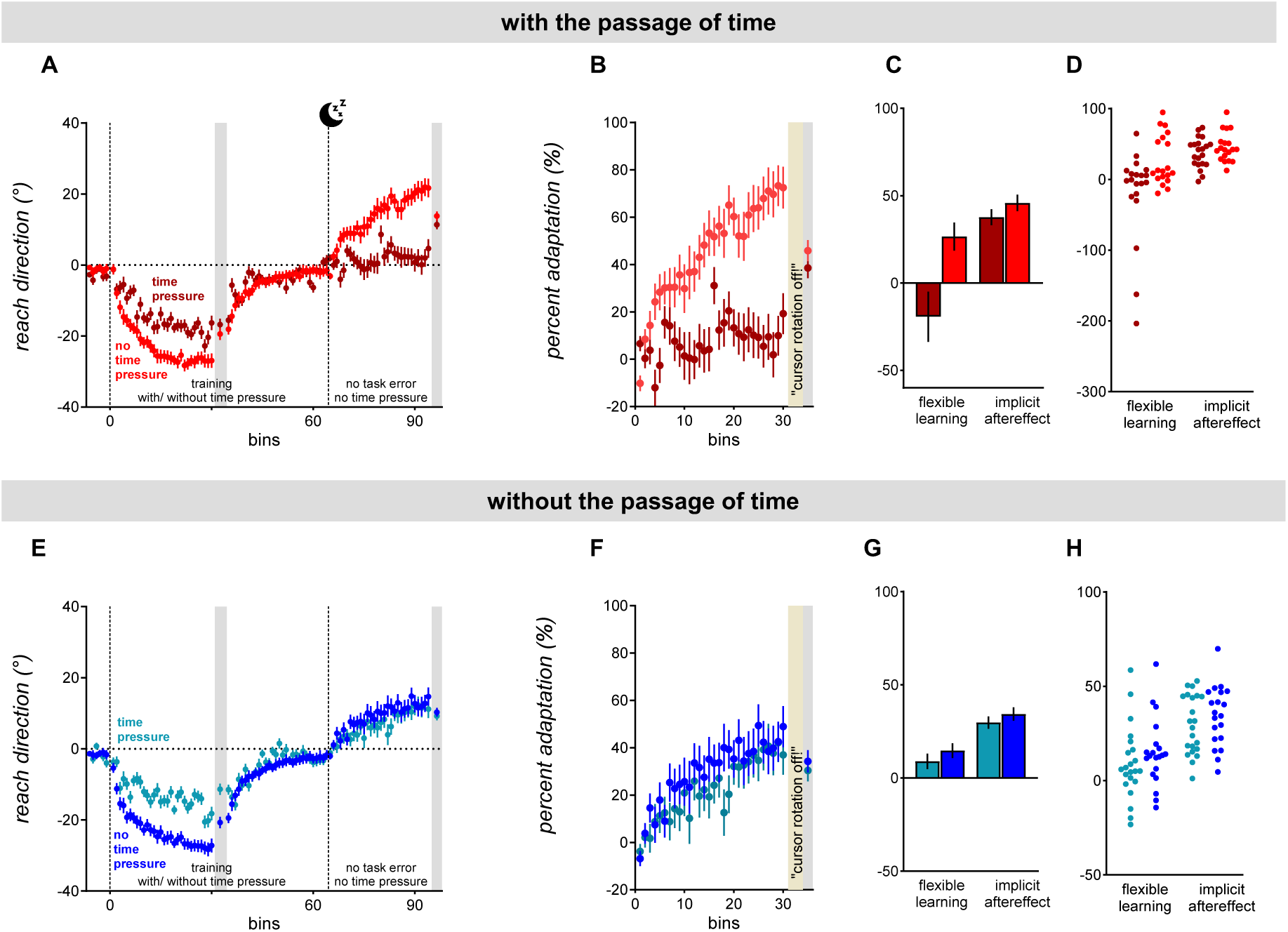
For those who had a time delay after training, previous training with time-pressure impaired subsequent adaptation to a conflicting perturbation even in the absence of task errors and time pressure. This time-sensitive improvement for those who previously experienced task errors co-occurred with improved explicit strategy use, quantified as the volitional disengagement of adapted movements upon instruction. Error bars are standard errors of the mean.

#### Flexible learning

The effect of time on flexible learning depended on whether time-pressure was applied during training [strong evidence for Time x Time-Pressure x Instruction interaction, BF_incl_ = 13.84, F(1,73) = 2.95, p = 0.09, ω² = 0.008]. Whilst those who trained without time pressure showed prominent flexible learning, both with the passage of time after training [F(1,73) = 10.86, p = 0.004] and without the passage of time after training [F(1,73) = 13.43, p = 0.002], those who trained with time pressure showed very variable and poorly adapted behaviour prior to instruction, particularly in those who also had the passage of time after training (see Figure 3B). Perhaps as a consequence of such variable behaviour prior to instruction, we did not see clear evidence for a change in adapted behaviour from pre-instruction 31.5%, 95%CI [18.5% 44.6] to post-instruction [34.6%, 95% CI [28.6% 40.6%] (moderate evidence for excluding the main effect of instruction, BF_excl_ = 12.19), both with the passage of time after training [BF_01,U_ = 3.26, F(1,73)=0.62 p = 0.44], and without the passage of time after training [BF_01, U_ = 5.06, F(1,73) = 2.21, p = 0.15].

#### Implicit aftereffect

The passage of time after training tended to increase the size of the block B implicit aftereffect, (main effect of time, BF_incl_ = 1.87, F(1,78) = 5.79, p = 0.02, ω² = 0.03; post-hoc t-tests: BF_10, U_ = 71.08, t(78) = 2.41, Cohen’s d = 0.29 [0.05 0.54]. We did not find clear evidence that prior training with time pressure altered the size of the implicit aftereffect, unequivocal evidence for including the main effect of time pressure, BF_incl_ = 0.44, F(1,78) = 2.46, p = 0.12, ω² = 0.009.

### Results summary

In Experiment 1, the passage of time affected a task-error driven component that *improved* adaptation performance and enhanced flexible learning component when faced with a conflicting perturbation. Previous work suggests that task errors trigger a component that requires considerable preparation time to re-aim movements and an inflexible component that can be produced at short reaction times (Leow *et al*., 2020). In Experiment 2, we tested whether time affects the fast process, by replicating the task-error conditions in Experiment 1, but additionally imposing time pressure during initial adaptation. This time, we found that the passage of time after training with time-pressure *impaired* subsequent adaptation when faced with a conflicting perturbation. Together, results from both experiments suggests that the passage of time exerts effects on dissociable processes that are triggered by task errors during learning: a time-consuming, flexible component, and a fast, inflexible component. Across both experiments, we also found effects of time on learning process(es) driven by sensory prediction errors, as indicated in post-adaptation implicit aftereffects: the passage of time after initial adaptation reduced anterograde interference, regardless of whether initial learning was acquired with and without task error history and with and without time-pressure during initial learning.

## Discussion

The question of how the passage of time alters memories resulting from sensorimotor adaptation is incompletely unresolved. Whilst several studies showed time-dependent decay of interference effects between opposing sensorimotor memories (Brashers-Krug *et al*., 1996; Shadmehr & Brashers-Krug, 1997; Shadmehr & Holcomb, 1997; Shadmehr *et al*., 1998), other studies showed no time-sensitivity (Bock *et al*., 2001; Goedert & Willingham, 2002; Caithness *et al*., 2004). Such discrepancies might be partly because adaptation is driven by mutually interactive processes (Miyamoto *et al*., 2020; Albert *et al*., 2022a), which could be differentially sensitive to the effects of time. Here, we re-examined the question of how the passage of time after initial sensorimotor learning affects subsequent adaptation to an opposing sensorimotor perturbation, using methods that help us dissociate components of adaptation driven by sensory prediction errors and components of adaptation driven by task errors. We found that time altered the interference effects of distinct learning processes encoded in the presence of task errors. Time caused a time-consuming, flexible learning process to improve the capacity to counteract an opposing perturbation, thereby markedly reducing anterograde interference. Using time-pressure manipulations at training to suppress this time-consuming, flexible component, we also found that time caused a fast, inflexible learning component to impair subsequent adaptation performance with an opposing perturbation. Time had a relatively subtle effect on the effects of learning driven by sensory prediction errors, reducing anterograde interference in implicit aftereffects resulting from adaptation to opposing sensorimotor maps. Our findings demonstrate parallel, distinct effects of the passage of time on the multiple components subserving adaptation.

### Time alters a time-consuming, strategic process driven by task errors

In Experiment 1, an overnight time delay improved subsequent adaptation performance when exposed to a conflicting perturbation, and this co-occurred with an increase in strategy use. The passage of time here thus appeared to enhance a strategic component of adaptation that is flexible enough to improve adaptation to an opposing perturbation. This finding corroborates work demonstrating that an overnight time delay can improve declarative memories (Jenkins & Dallenbach, 1924; Brown & Robertson, 2007), for a review, see (Dudai *et al*., 2015). This time- and sleep-dependent improvement of declarative learning is accompanied by a coupling between slow oscillations and sleep spindles during non-rapid eye-movement sleep (e.g., Barakat *et al*., 2011). Recent evidence shows that increased coupling between slow oscillations and spindles also occurs after visuomotor adaptation (Solano *et al*., 2024). Thus, it is possible that the benefit conferred by time here to the strategic component of adaptation relied partly on a period of sleep. Future research is required to clarify if a similar pattern of coupling between slow oscillations and spindles drive time-dependent benefits on strategic adaptation.

### Time alters a fast, non-strategic process driven by task errors

Previously, we showed that under typical conditions without time-pressure, task errors concurrently trigger a time-consuming, strategic process and a fast, non-strategic process (Leow *et al*., 2020). Here, in Experiment 2, to isolate effects of time on the fast, non-strategic process, we employed time-pressure at training, which prevented the execution of time-consuming explicit strategies. In contrast to Experiment 1, where time enhanced adaptation to an opposing perturbation, the passage of time after training *worsened* adaptation to an opposing perturbation in Experiment 2. Although we only employed time-pressure at training, the effects of this non-strategic component still manifested under test conditions without time pressure. Thus, time-pressure may not be required to unveil interference effects of this inflexible component. Other test conditions that suppress strategy use (such as an absence of task error) may be similarly effective. Our findings extend those of Albert *et al*. (2022b) who showed robust anterograde interference when successively learning opposing perturbations under time-pressure across all trials, and this anterograde interference was also altered by the passage of time.

How does the passage of time affect interference effects of the slow-flexible and fast-inflexible learning processes under typical learning conditions without time pressure? We speculate that although time alters the effects of both a fast, non-strategic process and a slow, strategic process, the strategic process might either mask expression of the non-strategic process, supress expression of the non-strategic process, and/or compensate for negative effects of the non-strategic process. Such tightly-coupled relationships between a strategic (and therefore explicit) component and non-strategic (and therefore implicit) processes of adaptation is consistent with recent computational modelling work demonstrating interactive relationships between distinct explicit and implicit task-error dependent components of adaptation (Miyamoto *et al*., 2020; Albert *et al*., 2022b), and previous experimental work demonstrating interactions between explicit and implicit processes during adaptation (Mazzoni & Krakauer, 2006; Keisler & Shadmehr, 2010; Taylor & Ivry, 2011). These findings of mutually competitive interactions between declarative and procedural memories are also consistent with other forms of learning, such as category learning (Ashby & Crossley, 2010) and motor sequence learning (Brown & Robertson, 2007).

### Time induces a decay of a component of adaptation driven by sensory prediction errors

Previous work shows that the passage of time after initial adaptation to one perturbation can reduce interference in implicit aftereffects when subsequently adapting to a conflicting perturbation, which showed reduced interference in aftereffects after longer 5 hour+ intervals than after minutes-long intervals (Brashers-Krug *et al*., 1996; Shadmehr & Brashers-Krug, 1997; Goedert & Willingham, 2002; Hamel *et al*., 2021). Our findings are consistent with this line of evidence, where we found that the passage of time reduced anterograde interference in implicit aftereffects resulting from adaptation to opposing sensorimotor maps. The effects shown here were relatively subtle compared to previous studies (Brashers-Krug *et al*., 1996; Shadmehr & Brashers-Krug, 1997; Hamel *et al*., 2021). This might have resulted from the late quantification of aftereffects here (after 240 trials of exposure to the opposing perturbation), in contrast to previous studies which quantified aftereffects more frequently throughout adaptation, and which found that the effects of time were primarily prominent upon initial exposure to the opposing perturbation (Shadmehr & Brashers-Krug, 1997). We interpret the effect of time here as a result of a decay of initial implicit adaptation to sensory prediction errors, such that initial learning interferes less with subsequent adaptation to an opposing sensorimotor map. The persistent anterograde interference in implicit aftereffects shown here that decays incompletely with an overnight delay is consistent with an implicit learning process that persists over delays of 60 seconds (Hadjiosif *et al*., 2023). This temporally persistent implicit process can result in worse-than-naïve adaptation, even when adapting to a previously encountered perturbation (sometimes termed “anti-savings”) (Leow *et al*., 2020; Avraham *et al*., 2021; Hadjiosif *et al*., 2023). However, this possibility requires systematic investigation in future research, as our study was not explicitly designed to quantify the contribution of temporally persistent versus temporally labile implicit learning processes to anterograde interference.

### A component of learning acquired under time-pressure, and its potential contributions to anterograde interference

Whilst our study focussed on the effects of time on the distinct components of learning, it also unveils previously unknown effects of a fast component of learning that does not require lengthy movement preparation times. Similarly, a learning component that is expressible under time pressure has previously been demonstrated (McDougle & Taylor, 2018). We show here that this fast process is expressed despite the removal of time pressure, and even under no-task-error conditions that disincentivise expression of task-error driven learning.

Exactly what constitutes this learning process? We speculate that this learning results from some association between features of the task context and the response (Dolan & Dayan, 2013). Time-pressure during training likely suppressed the time-consuming, flexible process, resulting in poorer adaptation performance as participants executed a fast, imperfect movement solution. We propose that execution of this fast, imperfect movement solution encouraged the formation of associations between the task context and the task response at training. Such associations might have been “cached” in working memory (McDougle & Taylor, 2018). When faced with a different context of a conflicting perturbation, persistence of these inflexible cached associations might have impaired subsequent adaptation to an opposing perturbation, even in the absence of time-pressure. Persistence of this inflexible learning might partly explain why force-field adaptation learning acquired under acute pain conditions continues to be expressed under pain-free conditions (Salomoni *et al*., 2019). Under typical adaptation conditions, effects of this fast, inflexible component might be masked by the time-consuming, flexible process, but we speculate that it can contribute to anterograde interference effects.

The idea that associative learning contributes to sensorimotor adaptation is not new (Shadmehr & Brashers-Krug, 1997; Shadmehr & Holcomb, 1997; Avraham *et al*., 2022). Anterograde interference has been attributed to a transient inability to inhibit a previously learned visuomotor association (Shadmehr & Brashers-Krug, 1997). Several previously documented features of anterograde interference are consistent with this idea. For example, in contrast to savings, anterograde interference is sensitive to the volume of movement repetition at training: the more a movement is repeated at training, the more persistent is the resultant anterograde interference (Sing & Smith, 2010; Leow *et al*., 2013; Leow *et al*., 2014; Leow *et al*., 2016; Hamel *et al*., 2021). Repeatedly pairing a response with the task context during training might help strengthen such associative memories, such that they are more rapidly retrieved when encountering an apparently similar task context (Leow *et al*., 2020). Similarly, interference is greater when there is more overlap between the appropriate action for initial and subsequent learning (Krakauer *et al*., 2006). The interpretation that inflexible associative memories contribute to anterograde interference might explain the previously surprising finding that protocols that should result in savings (i.e., where the same perturbation is encountered successively) fail to show savings when no-perturbation washout trials are experienced prior to subsequent learning (Hinder *et al*., 2007; Hamel *et al*., 2021). According to this interpretation, returning behaviour to the unadapted state via washout does not erase previously acquired context-response associations, but instils a new, competing context-response association (Villalta *et al*., 2015). Such associative memories seem susceptible to the effects of time, as suggested by findings that interfering effects of washout are weakened by a 24 hour delay after washout (Villalta *et al*., 2015).

The sensitivity of associative memories to subtle or overlooked task features might also have partly contributed to previous conflicting results with anterograde interference. In the original studies showing time-sensitive decay of anterograde interference effects, no-perturbation “catch” trials were interspersed into the training blocks, which might have made the resulting motor memories more susceptible to the effects of time by weakening the strength of the associative memories (Brashers-Krug *et al*., 1996; Shadmehr & Brashers-Krug, 1997; Overduin *et al*., 2006). In contrast, studies which did not show time-sensitivity did not include no-perturbation catch trials (Caithness *et al*., 2004; Miall *et al*., 2004; Cothros *et al*., 2006). For example, as discussed in (Overduin *et al*., 2006; Criscimagna-Hemminger & Shadmehr, 2008), the presence of no-perturbation trials interleaved into training might weaken associative motor memories, as more no-perturbation appear to reduce the size of interference effects (Overduin *et al*., 2006; Focke *et al*., 2013; Stockinger *et al*., 2014). Future studies are needed to test if the effects of time on the inflexible learning acquired under time-pressure here are sensitive to the effects of interleaved catch trials.

Whilst we propose that time-sensitive associative memories triggered by task-errors partly contribute to anterograde interference, such errors may not be *necessary* for anterograde interference effects. This is evident from our findings, where anterograde interference effects were prominent even without task-error-history (i.e., in the no-task-error groups in Experiment 1) and under conditions where task errors are minimal, such as when the perturbation is very gradually imposed (Leow *et al*., 2014; Leow *et al*., 2016). Moreover, anterograde interference effects are evident under paradigms without appreciable task or target errors, such as in standard force-field adaptation, where the hand hits the target despite the perturbation (Caithness *et al*., 2004; Cothros *et al*., 2006; Smith *et al*., 2006). Thus, associations between task or target errors and the adaptive response might not be necessary.

### Limitations and Future Directions

In this study, we did not systematically investigate other factors known to influence anterograde interference and its susceptibility to the effects of time. For example, repetition of the adapted movement at performance asymptote increases the amount of anterograde interference (Sing & Smith, 2010; Leow *et al*., 2014; Leow *et al*., 2016; Hamel *et al*., 2021). In paradigms quantifying retrograde interference in an ABA block structure, the amount of repetition at asymptote also renders initial learning resistant to retrograde interference effects (Krakauer *et al*., 2005). Another factor known to influence the effects of time on interference effects is the consistency of the perturbation: learning that occurs in the context of frequent no-perturbation trials interleaved appears more to susceptible to the effects (Criscimagna-Hemminger & Shadmehr, 2008). Future experiments should systematically investigate how the amount of movement repetition and the consistency of the perturbation influences time-sensitive anterograde interference effects shown in the components of adaptation driven by sensory prediction errors and task errors.

Another limitation is that in Experiment 2, failures to commence movement by the imperative signal resulted the trial being aborted (i.e., the screen was blanked and the trial was replaced at the end of each bin), which might result in some decay of temporally labile learning, as visuomotor adaptation learning shows a decay with time constants of 15-20 seconds (Hadjiosif & Smith, 2013; Hadjiosif *et al*., 2023). This presents a possible problem in the implementation of such time-pressure manipulations, and demonstrates a need for future experiments to explicitly investigate if decay results from such blanked-screen trials under task conditions with and without time pressure.

We also do not yet fully understand how the brain encodes the distinct adaptation components described, and how such motor memories evolve with the passage of time. Whilst previous work has described neural correlates of single-session adaptation to target errors and sensory prediction errors (Diedrichsen *et al*., 2005), we still do not fully understand how time alters the neural processes underpinning the components of learning driven by task error and sensory prediction errors. In force-field adaptation, where errors in performance might be perceived and processed differently to target errors in visuomotor adaptation (Ikegami *et al*., 2020), the extent of anterograde interference is associated with distinct patterns of activity in the ventral and dorsal basal ganglia and the prefrontal cortex (Shadmehr & Holcomb, 1997). Recent work has also demonstrated sleep-dependent components of visuomotor adaptation (Solano *et al*., 2024). Future studies which behaviourally dissociate the distinct adaptation components during neuroimaging techniques might help elucidate the neural mechanisms of these components, and the effect of time on these components of adaptation learning.

### Summary

Here, we demonstrate that various different components of movement adaptation to perturbations of sensory feedback are sensitive to the effects of time. Time enhanced the effect of a time-consuming component driven by task errors, possibly by enhancing strategic selection of actions that facilitate adaptation to an opposing perturbation requiring a novel movement solution. Time also altered an component of adaptation acquired under time-pressure, that resulted in greater impairment of adaptation to the opposing perturbation. Finally, time also altered a component of adaptation driven by sensory prediction errors, where an intervening time-delay after initial adaptation reduced interference resulting from subsequent acquisition of an opposing sensorimotor mapping. These effects of time on distinct components of sensorimotor adaptation shed new light on the long-standing question of how the passage of time alters motor memories. A better understanding of how time differentially alters distinct adaptation components will enable judicious design of training schedules in contexts involving motor learning, including operation of complex machinery, rehabilitation, and sport.

